# Olfactory Landmarks and Path Integration Converge to Form a Cognitive Spatial Map

**DOI:** 10.1101/752360

**Authors:** Walter M. Fischler, Narendra R. Joshi, Virginia Devi-Chou, Lacey J. Kitch, Mark J. Schnitzer, Larry F. Abbott, Richard Axel

## Abstract

The convergence of internal path integration with sensory information from external landmarks generates a cognitive spatial map in the hippocampus. We have recorded the activity of cells in CA1 during a virtual navigation task to examine how mice represent, recognize and employ sparse olfactory landmarks to estimate their location. We observe that the presence of odor landmarks at multiple locations in a virtual environment greatly enriches the place cell representation and dramatically improves navigation. Presentation of the same odor at different locations generates distinct place cell representations, indicating that path integration can disambiguate two identical cues on the basis of location. The enhanced place cell representation at one cue location led to the formation of place cells at locations beyond that cue and, ultimately recognition of a second odor cue as a distinct landmark. This suggests an iterative mechanism for extending place cell representations into unknown territory. These results reveal how odor cues can serve as landmarks to guide navigation and suggest a model to explain how the convergence of landmarks and path integration participates in an iterative process that generates a cognitive spatial map.

---

Hippocampal representations of an animal’s environment reflect both external sensory landmarks and internal path integration of the animal’s movement in space^1^. These two sources of a cognitive spatial map, path integration and landmarks, are mutually dependent^2–7^. Path integration uses idiothetic signals generated by self-motion to define an organism’s position relative to fixed points or landmarks^8–10^. Such landmarks are valuable only if they are recognized as fixed in space, and this determination may require path integration^11^. Sensory landmarks and internal path integration signals are likely to converge in the hippocampus. We examined hippocampal activity in mice performing a navigational task that relies solely on path integration and sparse olfactory cues. These observations provide a new model that explains how the convergence of sensory and idiothetic signals drive the development of a hippocampal spatial map. Our results suggest that the extension of a spatial map into previously unexplored territory is an iterative process in which path integration from existing landmarks identifies new landmarks that then provide a basis for further path integration.

Olfactory cues are a primary source of sensory information in mice and can serve as landmarks when fixed in space. The hippocampus receives olfactory information from the lateral entorhinal cortex (LEC)^12–15^. The LEC receives input directly from the olfactory bulb and piriform cortex^16,17^, two structures that encode odor identity. The influence of odors on hippocampal activity has been shown in both spatial and non-spatial contexts^18–21^. Grid^22^, head direction^23^, boundary^24^, and speed cells^25^ in the medial entorhinal cortex (MEC)^26^ provide information to the hippocampus about location and self-motion in real and virtual environments^27^. The hippocampus thus has access to both the position and identity of an olfactory landmark.

We created a virtual spatial navigation task that required mice to combine path integration signals with olfactory cues, which provided the only sensory landmarks. While mice performed this odor-guided virtual navigation task we monitored large-scale neural ensemble activity in the CA1 region of hippocampus using a head-mounted miniature fluorescence microscope^28,29^. Head-fixed mice ran on a featureless spherical treadmill in total darkness and received a water reward after they traversed a virtual linear distance of 4m from the starting point. The ball had a single rotational axis, rendering the task equivalent to navigation on a linear track. Mice first performed this task without sensory cues for 1-2 weeks (1 session per day, ≥ 60 trials per session). We then introduced 1s odor pulses delivered when the mice reached locations 1m and 3m from the starting point (figure 1a). The same odor was used at both locations in a given trial, but trials with two different odors (limonene and pinene) were interleaved randomly. This task required the mice to determine their location solely on the basis of path integration plus minimal olfactory cues. Furthermore, because the same odor was introduced at two locations, the mice had to use path integration to disambiguate the location of the two odor cues. This experimental paradigm permitted the study of the convergence of idiothetic and external olfactory information in the generation of cognitive spatial maps.

**Figure 1.**
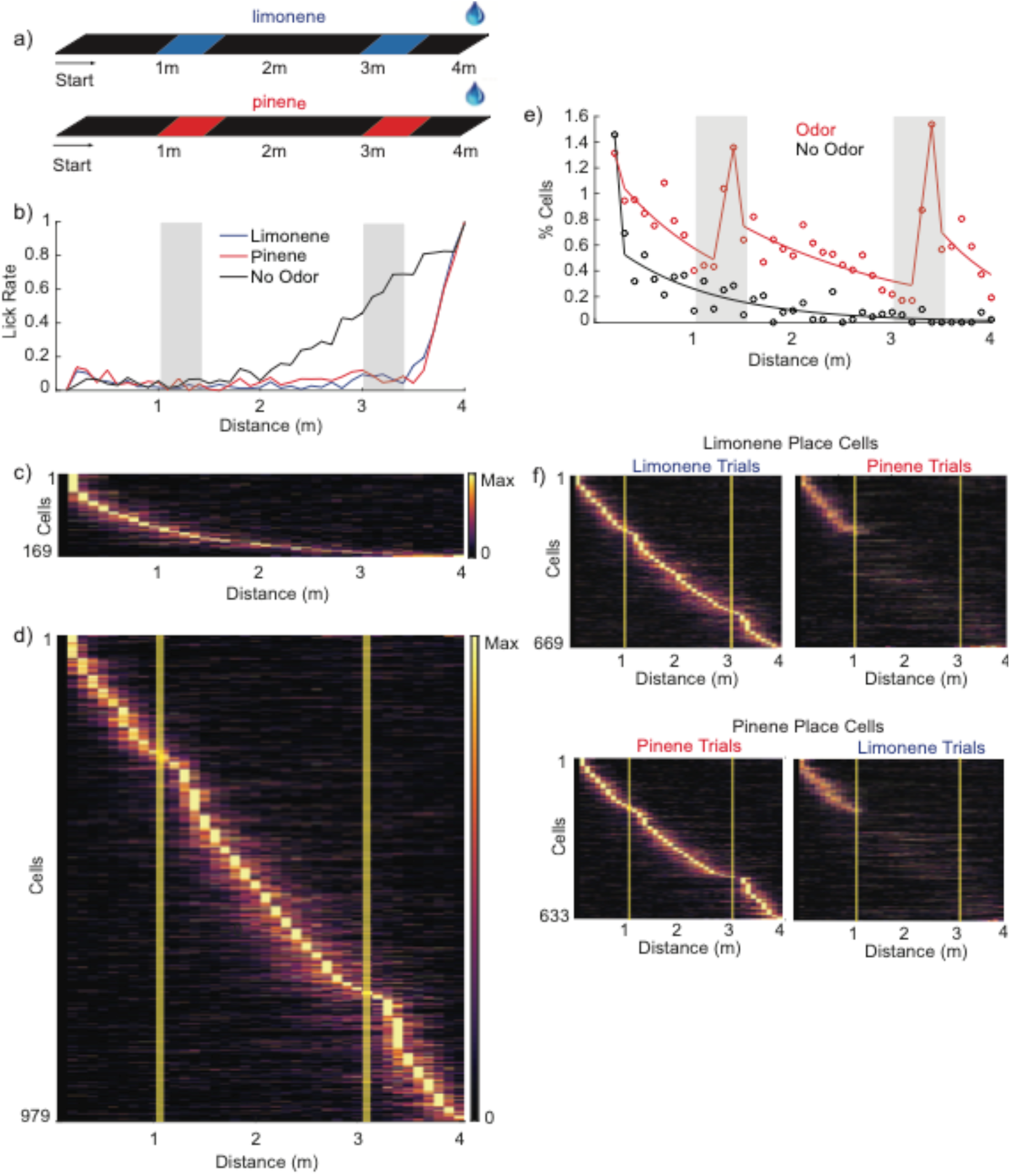
Odor landmarks dramatically enhance place cell representations and improve navigation behavior. **a**, Schematic of the virtual track with odor landmarks. Either limonene (blue) or pinene (red) is presented for 1s at both 1m and 3m depending on trial type. **b**, Mean rate of licking at positions along the track for different trial types (n = 5). Last session with no odor landmarks (black) and on 4th day with odor landmarks, limonene trials (blue) and pinene trials (red). Shaded grey areas denote 1s odor pulses on odor trials. **c, d**, Activity of combined place cells for all mice (n = 5) along the virtual track sorted by the position of peak mean activity. **c**, Activity on last session with no odor landmarks (6%, 169/2893 cells). **d**, Activity on 4th day with odor landmarks (35%, 979/2778). Yellow lines denote onset of 1s odor pulses. **e**, Distribution of place fields across the virtual track. Mean density of place cells, as a percentage of all recorded cells (n = 5). Black circles, last session with no odor landmarks. Black line, exponential fit to the data between the peak at the start and 4m. Red circles, trials on 4th day with odor landmarks. Red line, exponential fits to the data from peak at the start and 1m odor landmark, between the 1m and 3m odor landmarks, and 3m odor landmark to 4m. **f**, Activity of place cells on 4^th^ day of training with odor landmarks on either limonene or pinene trials. Top, cells sorted by place fields on limonene trials, activity on limonene trials (left) and pinene trials (right). Bottom, cells sorted by place fields on pinene trials, activity on pinene trials (left) and limonene trials (right).

After performing the task in the absence of odor cues for 1-2 weeks, the mice initiated anticipatory licking and decreased their running speed after traveling about 2m along the 4m track (figure 1b and supp. figure 1a). This premature licking 2m before the reward location suggests that path integration alone cannot accurately measure distances greater than 2m. The mice then performed the task in the presence of odor cues at 1m and 3m locations for 4 days, after which the animals suppressed licking and maintained high running speeds for ~3.5m of travel, licking only ~0.5m before the goal location (figure 1b and supp. figure 1a). This suggests that the mice recognized the odors as spatial landmarks and used these landmarks to improve navigation.

We used a miniature microscope (nVista 2.0, Inscopix) and the genetically encoded fluorescent Ca^2+^ indicator GCamp6f to image the somatic Ca^2+^ activity of ~2,400–3,000 CA1 pyramidal neurons per session in 5 mice. We computationally extracted individual neurons and their Ca^2+^ activity traces from the fluorescence videos and registered their activity patterns to the trajectories of the mice on the virtual track (see methods). Neurons with consistent position-selective activity were classified as place cells (see methods).

After 1-2 weeks of training without odor cues, 6% (169/2893 total cells) of the imaged neurons exhibited the properties of place cells (figure 1c). The number of place cells was maximal at the starting location and decreased exponentially with distance along the track (figure 1e), 86% (146/169) of place cells had place fields at locations less than 2m from the start. In addition, the variability in the activity of recorded cells across trials increased with distance from the start location (supp. figure 2a,b). These results suggest that path integration alone can only support the formation and activity of place cells over distances less than 2m. The sparsity of place cells beyond 2m is consistent with the behavioral observation that mice could not reliably determine their location beyond this distance.

After mice performed the task for 4 days with odor cues at 1m and 3m we observed a 6-fold increase in the percentages of neurons that qualify as place cells. In trials with either limonene or pinene 35% of recorded cells were classified as place cells (979/2778 total cells) (figure 1d). This increase occurred at all locations but was particularly pronounced at 1m and 3m, the locations of odor exposure. Local peaks in place cell density appeared at the start of the virtual track and at the sites of each of the two odor cues (figure 1e). Between these local peaks, the number of place cells decreased exponentially as a function of distance. The variability in the activity of recorded cells across trials also increased as a function of distance from the start location and both odor cues (supp. figure 2a,b). The presence of three spaced peaks caused the place cell density to remain high along the entire track despite the exponential decline between the peaks. The elevated place cell density correlated with the animal’s ability to suppress licking and retain running speed up to the reward site when odor cues are present. These results demonstrate that brief olfactory cues create landmarks that can combine with path integration to generate a robust ensemble of place cells in support of accurate navigational behavior.

Performing the same experiment with two different odors allowed us to examine the generation of spatial maps in two different sensory contexts. The two sensory contexts generated different place cell representations, demonstrating remapping. As expected, place cells between the start site and the first odor cue were the same in the two odor contexts (figure 1f). However, following the first exposure to odor, the limonene and pinene trials evoked different sets of place cells. Only 11% (78/708) of all place cells with fields beyond 1m, the location of the first odor cue, were shared at the same locations in the two contexts. In accord with these findings, population vectors (PV) representing the neural ensemble activity during the two trial types were highly correlated for the first 1m of travel, but these correlations decreased to ~15% after the onset of the first odor and remained low at all positions beyond 1m (supp. figure 3). These data indicate that different odors generate distinct cognitive spatial maps under conditions with identical task contingencies.

We next asked whether the continued presence of odor landmarks is needed to maintain the place cell map and the associated improvement in behavior. After 4 days of experience with odor cues, the mice performed additional sessions in which odors were presented on only half of the trials. The density of place cells in trials without odor cues exhibited a single peak at the start location in contrast to the three peaks observed when odor cues were present. Beyond the start location the density of place cells decayed exponentially with a length constant of 1.5m (figure 2d), only slightly greater than the 1m length constant seen prior to odor training. Moreover, the number of place cells beyond 2m was 2.4 times greater on trials with odor cues than on trials with without odor cues, 308 and 128 respectively (figure 2a, 2b). In accord with the neural activity, anticipatory licking and slowing were less accurate on trials without odor cues than on trials with cues (figure 2c, supp. figure 1b). These observations suggest that the generation of a robust place cell map and navigation behavior are enhanced by the continued presence of odor landmarks.

**Figure 2.**
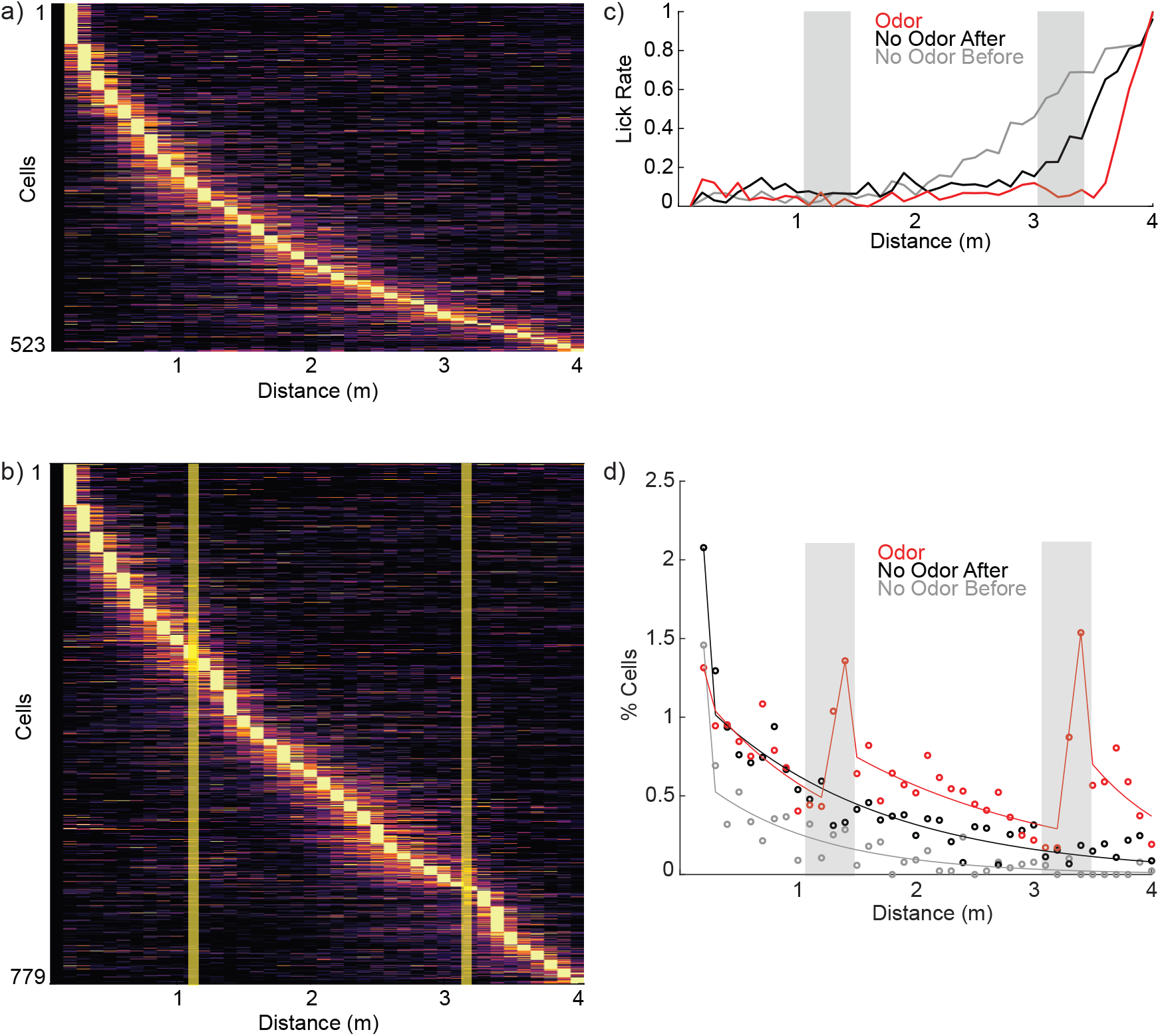
Odor landmarks are required to enhance place cell representations. **a**, Activity of place cells on trials with no odor landmarks after odor training (16.9%, 523/3087 cells). **b**, Activity of place cells on trials with pinene odor landmarks after 4 days of training (25.2%, 779/3087). **c**, Mean rate of licking at positions along the track for different trial types (n = 5). Grey, last session with no odor landmarks before odor training. Red, after 4 days with odor landmarks. Black, 4 days of odor training followed by one day without odor. **d**, Distribution of place fields across the virtual track. Mean density of place cells, as a percentage of all recorded cells (n = 5). Colors as in **c.** Curves are exponential fits as in **figure 1e**.

We next explored the dependence of place cell activity on reward. After 5 days of training in the presence of odor cues, the mice performed an additional session in which the reward site was absent. Under these conditions the mice ran at similar speeds but did not lick or stop at the previously rewarded site, and the percentage of place cells decreased from 35% to 1% (28/3919 cells) (figure 3c). Thus, the activation of a robust place cell representation is contingent on both odor landmarks and reward.

**Figure 3.**
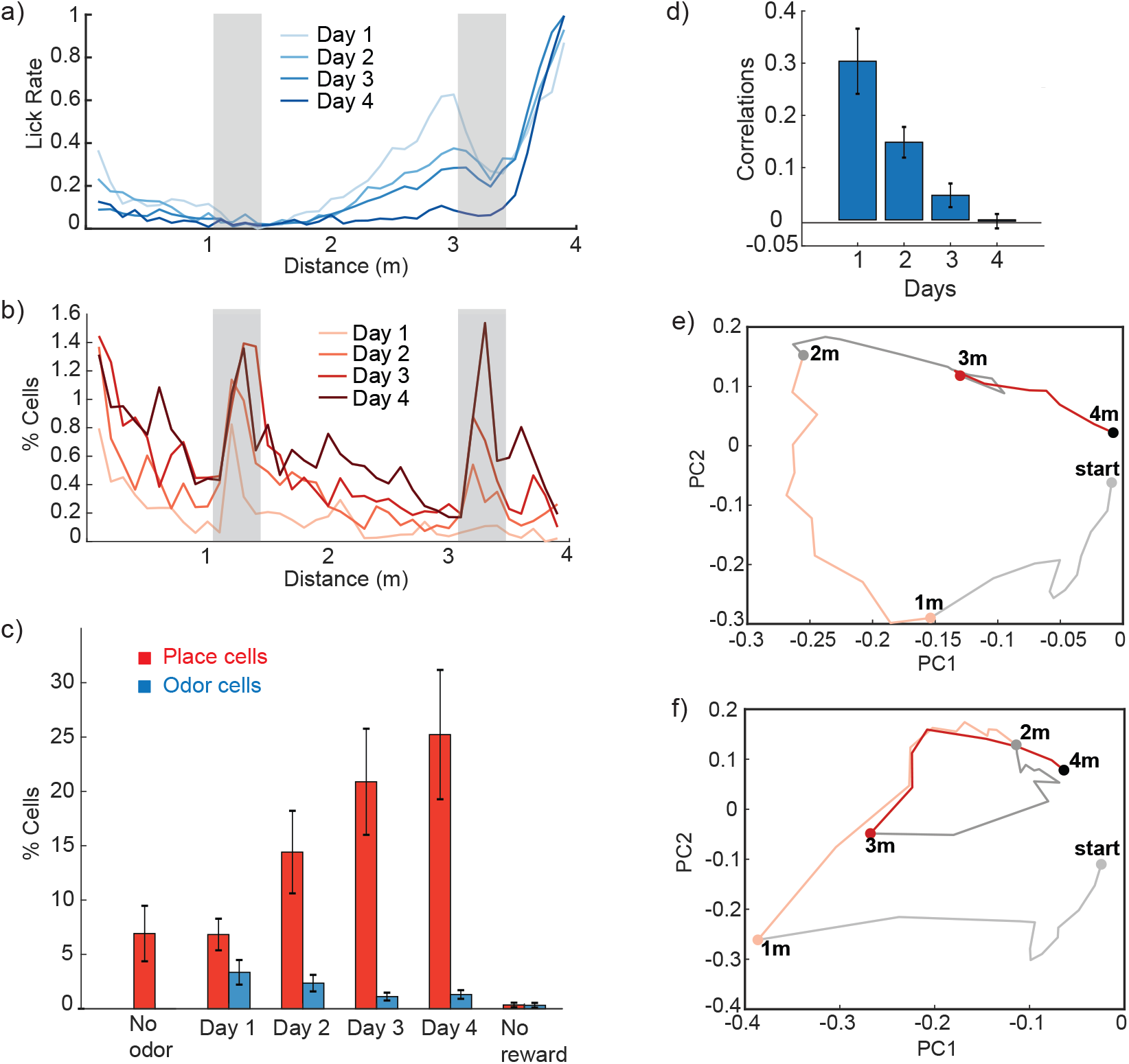
Evolution of place cell maps and improvement in navigation behavior. **a**, Mean rate of licking at positions along the track on each day (n = 5). **b**, Distribution of place fields across the virtual track. Mean density of place cells, as a percentage of all recorded cells, at positions along the track on each day (n = 5). **c**, Mean percentage of place cells (red) and odor cells (blue) on each day (n = 5, error bars = SEM). **d**, Mean Pearson correlation per mouse (n =5, error bars = SEM) of the activity population vector of all recorded cells from 1m to 2m and 3m to 4m. **e, f**, Population vector trajectory along the virtual track on pinene trials in the 2-dimensional space defined by the first 2 principal components for all recorded cells. **e**, 4 days of training. **f**, 1 day of training.

We further analyzed the interplay of path integration and landmarks by examining the emergence of place cells during training and its relation to improved navigational behavior. On the first day of training in the presence of odor mice initiated licking and decreased their speed at ~2m as observed before odors were introduced. However, the mice transiently reduced their lick rate at the 3m odor cue, suggesting that they had learned that the odor cue is distant from the reward site (figure 3a). Over the next 3 days of training, we observed a gradual reduction of licking at locations that preceded reward and by the fourth day of training the mice avoided licking and maintain a high running speed up to 3.5m (figure 3a, supp. figure 1c).

The evolution of the place cell representation correlated well with the successive improvement in navigational behavior. On the first day of training with odor cues we observed the emergence of a large number of odor cells, neurons that responded similarly at the two sites of odor presentation (figure 3c, supp. figure 4b). In addition, cells with place fields at the 1m odor cue increased in number, whereas cells with place fields at the 3m odor cue largely remained absent (figure 3b, supp. figure 4a). Over the subsequent days of training, the percentage of place cells steadily increased, whereas the number of odor cells steadily decreased (figure 3c, supp. figure 4c). Importantly, the emergence of a peak in the density of place cells at 1m was accompanied by an increase in the percentage of place cells between 1m and 3m. Over the course of several days, as more place cells formed in the region between 1m and 3m, an additional peak in place cell density arose at 3m (figure 3b). The emergence of peaks in the place cell densities at 1m and 3m was thus gradual and sequential. This suggests that an increased number of place cells at one location can enhance the formation of place fields further along the animal’s trajectory. This results in the emergence of place cells beyond 3m, coinciding with the improvement in behavior.

These results suggest a sequence in which first, the odor cue nearest the start location is recognized as a landmark, resulting in a peak in place cell density at 1m. This recognition then leads to a gradual increase in place cell density at locations between 1m and 3m. The extension of the place cell map to 3m then allows the animals to recognize the 2^nd^ odor cue as a different landmark, resulting in a second peak in place cell density at 3m. Recognition of this second landmark supports the creation of additional place cells beyond 3m, and this leads to more precise navigation to the goal. This sequential iterative process may be a basic mechanism for extending place cell representations into unknown territory.

We complemented the study of individual place cells with an analysis of population level activity. We used a population vector to represent the activity patterns of all imaged neurons and compared the correlations in the population vectors at the two locations where odor cues were presented. These population vectors were strongly correlated on the initial day of odor training, but this correlation gradually declined to near zero over the course of training (figure 3d). This decline parallels the transformation of odor cells into place cells at the locations of the odor cues and reflects, at the population level, the recognition of distinct spatial landmarks.

We performed a principal component decomposition to study the dynamic trajectories of the neural ensemble activity across all imaged cells. A projection of the trajectory of the population activity onto the first two principal component vectors shows an interesting relationship to the task. This virtual task has the topology of a circle because the mice ‘return’ to the start position on the next trial after reaching the reward location. After 4 days of odor training, when the animals have developed an accurate sense of the location of the odor cues and reward, the PC-projected population activity has a circular shape and the locations of the odor cues are appropriately spaced along this trajectory. (figure 3e) Thus, the trajectory of activity in neural ‘state’ space bears a striking topological and metrical similarity to the virtual space of the task.

We also examined the evolution of the state-space trajectory during odor training. On the first day of odor exposure, the state-space trajectory corresponding to locations near 2m is close to the point on the trajectory representing the reward location. When the 3m odor cue is delivered the trajectory loops back to the location of the 1m odor cue and then closely follows its previous path (figure 3f). On the second day of training, backward looping is reduced and by the 4th day the points of the state-space trajectory near 2m are well separated from the representation of the reward site (supp. figure 5a,b). These results are consistent with a transformation of the internal representation of odor cues into two distinct odor landmarks that serve to guide navigational behavior.

Our experiments reveal that odor cues can serve as landmarks to guide navigation, and they suggest a model to explain how the convergence of landmarks and path integration generates a cognitive spatial map in the hippocampus. The model we constructed consists of a population of place cells driven by inputs from a set of path integrators, and feedback from the place cells back to the path integrators (figure 4a). In the absence of odor cues, each path integrator generates an independent estimate of the distance that the animal has travelled from the starting point, and each estimate drives a different spatially modulated input to the place cells. The path integrators are noisy so their estimates of position and, consequently, the inputs they drive vary from trial to trial. The trial-to-trial variance grows with the distance travelled as path integration becomes less reliable (figure 4b).

**Figure 4.**
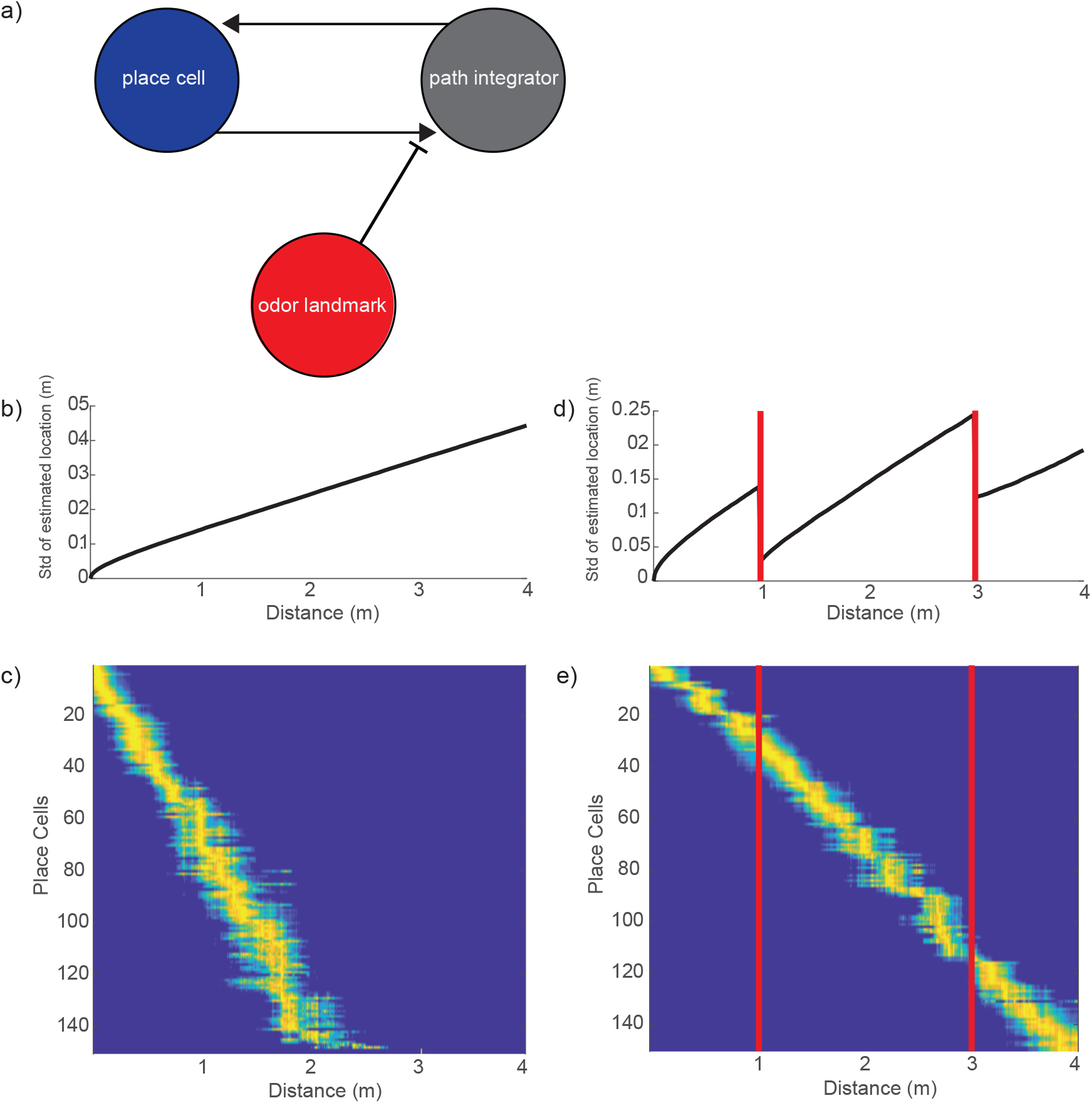
A model for the convergence of path integration and odor landmarks in place cell formation. **a**, Schematic of the 3 components of the model with reciprocal connections between place cells and path integrators. **b**, The trial-to-trial standard deviation of position estimates by a population of model path integrators in the absence of odor landmarks. The standard deviation and hence positional uncertainty grow monotonically with distance travelled. **c**, Model place cells formed in the absence of odor landmarks **d**, The trial-to-trial standard deviation of position estimates by the population of model path integrators in the presence of odor landmarks. The standard deviation decreases at the locations of the odor cues due to resetting of the path integrators by the place cells. Red lines denote locations of odor input. **e**, Model place cells formed in the presence of odor landmarks. Red lines denote locations of odor input.

In the model, place cells form by a process that simulates the effects of plateau potentials^30,31^. The model suggests that, in each cell, a plateau potential occurs at a random location, resulting in plasticity that sets the weights of the synapses from the integrators to the place cell to values proportional to the presynaptic input at the time of the plateau. Following this plasticity, the cell performs template matching, responding if there is a close enough match between the current input rates and the rates experienced at the time of the plateau. This process creates reliable place cells at short distances from the starting location because the inputs driven by path integration are similar from trial to trial at these locations and therefore well matched to the template. For large distances, on the other hand, inputs vary considerably from trial to trial and rarely match the template. As a result, reliable place cells cannot form. We chose a level of noise for the path integrators so that reliable place cells form at a distance less than 2m but not at greater distances (figure 4c). In agreement with the model, the dominant contributor to the variability of place cell responses in the data (supp. figure 2a) is failure of place cells to fire reliably across trials (supp. figure 2b).

Model place cells project back to the path integrators, and this pathway is also subject to plasticity^7^. Thus, at the same time that plasticity modifies synapses from path integrators to a place cell, it also modifies connections from this place cell back to the path integrators. This plasticity stores a trace of the distance estimate provided by each path integrator at the time of the plateau. The result is a pathway by which the activity of a place cell can reset the path integrator^7^ back to the value it took when the place field formed. In our model, this pathway is only engaged when a sensory cue, the odor, appears.

In the model, when an odor appears, place cell activity drives the path integrators to their previously stored values (figure 4d). Although these values are no more accurate than the estimates of distance on any other trial, they are consistent from trial to trial due to the reset provided by the odor-activated place cells. Thus, place cells that form beyond the 2m point have inputs that are more reliable as a consequence of odor landmarks. This consistency allows reliable place cells to be created by the plateau potential mechanism. This results in a generative process that produces a complete place cell representation along the entire 4m track (figure 4e).

The system we have described consists of two networks, the place cells and the path integrators, that store within their synapses the traces of their relationship at the time of place cell formation. Place cells are maximally driven by path integrators that match the input when their place fields formed. Reciprocally, place cell inputs to each path integrator store the value of that path integrator when the place field formed. This system is calibrated by an external event that identifies when these relationships are consistent. This event is a landmark.

Our experiments are in accord with several features of the model. The same sensory information at different locations can generate distinct place cell representations. Thus, path integration can disambiguate two identical cues on the basis of location. Moreover, different odor cues at the same location generate different place cell representations that extend beyond the odor cue. These observations are consistent with the role of the hippocampus in the transformation of egocentric sensory information into allocentric cognitive spatial maps of the external world.

In our experiments the enhanced place cell representations at the location of a brief odor cue led to the formation of place cells at locations beyond that cue. This implies that the number of place cells active at one location influences the number of place cells active at subsequent locations. In the absence of new information from the senses, this influence diminishes as the path integrator becomes progressively less accurate. Place cell densities show a quantitatively similar exponential decrease as a function of distance either from the start location or from the location of olfactory landmarks. Thus, the presence of olfactory cues appears to reset the path integrator, as suggested by our model.

The ability of an odor cue to serve as a spatial landmark depends on the accuracy of the path integrator at positions leading up to the odor location. When the same odor cue is present at two distinct locations the cue nearest the starting position (itself a landmark) is first to generate a unique place cell representation and appears to reset the path integrator. Over the course of training, place cells are generated that span the gap between the two spaced, but identical, odor cues. Thus, identical sensory features reliably present at multiple locations can be identified as unique landmarks by means of a generative process that relies on path integration. The addition of new landmarks could then further extend the cognitive spatial map. This allows the convergence of path integration signals with landmarks in the hippocampus to construct a spatial map that supports navigation over distances far greater than path integration alone.

## Author contributions

W.M.F., R.A., L.F.A, and M.J.S. designed experiments and analysis. W.M.F., N.R.J., and V.D. performed the experiments. L.J.K. and M.J.S. designed a calcium signal extraction algorithm used for the initial analysis of the imaging data. L.F.A. generated the computational model. W.M.F., R.A., L.F.A., and M.J.S. wrote the paper, and all authors commented on the manuscript.

## Acknowledgments

We thank Attila Losonczy, Matt Lovett-Barron, and Patrick Kaifosh for experimental advice. Bert Vancura and Yolanda Zafrina for technical assistance. Dmitriy Aronov for comments on the manuscript. Barbara Noro for helpful comments and suggestions. Clayton Eccard for assistance in preparation of the manuscript. Phyllis Kisloff, Miriam Gutierrez and Adriana Nemes for general laboratory support. This work is supported by The Helen Hay Whitney Foundation (W.F.), NSF NeuroNex Award DBI-1707398 (L.F.A.), the Gatsby Charitable Foundation (L.F.A.), the Simons Collaboration for the Global Brain (L.F.A.), and Howard Hughes Medical Institute (M.J.S. and R.A.). R.A. and M.J.S. are HHMI investigators.

## Methods

### Animals and surgery

All experiments were approved by the Columbia University Institutional Animal Care and Use Committee. Adult male C57BL/6 mice (aged 8-12 weeks) underwent two surgical procedures under isoflurane (1-2%, vol/vol). We injected ~500nl of a 1:3 dilution in PBS of UPenn Vector Core packaged AAV2/1 serotype virus expressing GCaMP6f under the control of the CaMKII promoter (AAV1.CamKII.GCaMP6f.WPRE.SV40, titer 1-3 × 10^13^ vg/ml) with a thin glass pipette into the left hemisphere of dorsal CA1 (−2.2mm from bregma, 1.6mm mediolateral, −1.2mm dorsoventral). 1-2 weeks after viral injection we implanted a 1.8mm diameter imaging cannula (metal cannula with a glass coverslip attached at the bottom, Inscopix part #: 1050-002189) over the dorsal surface of CA1 centered on the site of viral injection after aspiration of the overlying cortical area as previously described^32^. We then secured the cannula and a custom metal head bar to the cranium of the mice using dental cement (Dentsply). 1-2 weeks after cannula implant we inserted a 1mm diameter gradient refractory index (GRIN) micro-endoscope (Inscopix part #: 1050-=002176) into the cannula and a plastic baseplate (Inscopix part #: 1050-002192) was cemented into place after confirming even expression of GCaMP6f in healthy tissue using a miniaturized fluorescent microscope (Inscopix nVista, v2.0).

### Virtual odor-guided navigation system

Mice were head-fixed on a spherical treadmill (20cm diameter Styrofoam ball) rotating on a single axis. The axis of the treadmill was attached to an analog rotary encoder (US Digital part #: MA3-A10-125-B) connected to an Arduino Mega2560. Angular displacement was converted into a virtual linear distance based on the circumference of the treadmill. A water port consisting of a small gavage needle (Cadence Science part #: 7901) connected to a water reservoir was placed within reach of the mouse’s tongue. A capacitance touch sensor (Sparkfun #MPR121) was attached to the water port to measure licking and the sensor was connected to the Arduino Mega2560. Small 2-4ul drops of water were delivered by the brief opening a solenoid valve (Lee Valves #LHDA 12712154) connected to the water port. Custom Arduino software was used to deliver water drops at reward locations. Limonene and pinene odor cues were delivered via a custom olfactometer controlled by an Arduino Mega2560. 10% solutions of limonene (Sigma #183164) and pinene (Sigma #P45680) diluted in mineral oil (Fisher Scientific #0121-1) were added to syringe filters (Whatman #6888-2527) and an additional filter of pure mineral oil was used to provide blank odor stimuli between the 1s presentations of limonene and pinene cues. Custom Arduino software was used to control odor valves for switching between limonene or pinene and blank (mineral oil) filters. Two mass flow controllers (MFC) were used to maintain a constant airflow of compressed medical grade air for odor delivery. One MFC was set to deliver air to the odor and blank filters at 0.3 L/min. The other MFC was set at 0.7 L/min to deliver clean air for a carrier stream. The combined airflow experienced by the mouse was a constant 1 L/min in the absence or presence of limonene and pinene odor cues. The odor or blank air streams and the carrier stream were combined in an 8-port odor manifold (Island Motion Corporation 020206.0001) connected to one side of a custom odor port that was placed within 2mm of the nose. A vacuum was connected to the opposite side of the odor port. The vacuum line was controlled by an MFC set at 1 L/min to remove air and odor continuously from the odor port. Speakers delivering white noise at 70 dBs were placed in front of the mouse to cancel out ambient noise and the sound of the valves opening and closing. The entire experimental system was enclosed by black hardboard (Thorlabs TB4) on the sides, Blackout nylon fabric (Thorlabs BK5) on the top, and the lights were kept off in the room to maintain a dark environment. Mice were monitored using an IR camera (Basler A601f) and illuminated using an IR light.

### Behavior training

After surgeries, mice were place on a 12-hour reverse light/dark cycle. All experiments were performed in the middle of the active (dark) period. Mice were habituated to handling for several days. Mice were then habituated to head-fixation on the spherical treadmill for several days before being put on water restriction. After 2-3 days on water restriction (~2 ml water per day), mice were then trained to walk increasing linear distances to receive water rewards. Water rewards consisted of 2-4ul water drops triggered by 2 consecutive licks over 4s. The beginning of the water reward epoch was signaled by the emergence of a single 2-4ul drop on the lick port. At the end of the 4s water reward window the virtual distance was reset to 0 and the next trial began. Initial distance to reward was set at 0.5m. After mice were able to complete >60 trials in one 20min session (1 session/day) the distance was gradually increased from 0.5 to 1m and then from 1m to 4m in 1m increments. Upon beginning of training on the 4m virtual track, the area in which water rewards were available was restricted to a spatial window from 4m to 5m. At the 4m distance mice were required to complete >80 trials in a single 20 min session for 3 consecutive days, at which point we began collecting the data for these experiments. The 5 mice used in this study reached criteria after 5-10 days of training at 4m.

### Imaging and behavior data collection

At the beginning of each experimental session, mice were head fixed on the spherical treadmill and the miniature microscope (Inscopix, v2.0) was attached to the plastic baseplate. The field of view containing G-Camp6f expressing neurons was examined to confirm that the site was aligned with previous recording sessions. Imaging data was collected at a frame rate of 20Hz. LED power was set between 30-40%. The data was initially collected at a resolution of 1440 × 1080 pixels and then subsequently down-sampled by a factor of 4 for further analysis.

### Processing of imaging data

Calcium imaging movies were preprocessed using the Mosaic software package (Inscopix). During preprocessing, movies were spatially cropped to fit the imaging site and motion corrected. Individual neurons were isolated using published CNMF-E Matlab code^33^. The output of the deconvolved signal *S* based on the OASIS algorithm^34^ for each identified cell was used for further analysis of neural activity.

### Behavioral data processing and alignment to neural activity

We used custom Matlab software to convert rotary encoder signals to a virtual linear distance and speed was calculated over rolling 200ms time windows. Lick detection from touch sensor signals was aligned to virtual distances. A TTL pulse was sent to the Arduino Mega2560 from the microscope on the acquisition of each imaging frame to align neural activity to virtual distance, speed, and licking. Signals for the opening and closing of odor and water reward valves were recorded by Arduino Mega2650 and aligned to behavior and activity data.

### Spatial binning and selection of trials for analysis

Aligned behavioral and imaging data was averaged within spatial bins of 100mm. We excluded the first spatial bin from 0-100mm and any data after initiation of the initial water reward to limit our analysis to periods when the mice were actively running towards the next goal location. The first 5 trials for each trial type in a session were excluded as there was no explicit signal for the start of the experiment and several 4m reward crossings were required before mice showed awareness of the task. We analyzed trials 5-30 for each trial type so as to confine the study to epochs with constant running (speed > 5cm/s) along the virtual track and similar levels of motivation. Mean lick rate and speed over all trials in a session was normalized to the max lick rate and speed for each individual mouse and then averaged across all 5 mice. Data on limonene and pinene trial types was averaged for each mouse on all plots except figure 1b and S1a.

### Place cell analysis

We classified neurons as place cells after meeting three criteria: 1) Activity was averaged over all trials and the bin with peak mean activity had > 3 times the mean activity over all bins. 2) Activity on all trials was z-scored and cell had z > 1 in the bin of peak mean activity on > 25% of all trials of that trial type 3) The virtual track was divided into 4 equal sectors (0-1m, 1-2m, 2-3m, and 3-4m) and criteria both 1 and 2 were met on one and only one sector of the track. Additionally, we classified neurons as odor cells if both criteria 1 and 2 were met on both sectors of the track in which odor cues were present (1-2m and 3-4m). Mean activity over all trials was normalized to the max mean activity for each neuron.

### Population vector (PV) activity

The variability in neural activity from trial to trial across the virtual track was calculated by using Pearson’s correlation on the spatially binned PV activity (mean for all trials, combination of all recorded neurons in all 5 mice) on odd and even numbered trials. Comparison of the activity at odor cue areas (1-2m and 3-4m) was calculated using Pearson’s correlation of the spatially binned PV activity (mean for all trials) of all recorded cells in individual mice and taking the average of all 5 mice. Comparison of activity between limonene and pinene sensory contexts calculated using Pearson’s correlation of spatially binned PV (mean over all trials) of all recorded cells in individual mice and taking mean of all 5 mice.

### Principal component analysis

To calculate the population trajectory, we applied the Matlab PCA algorithm to the spatially binned PV activity (mean for all trials) on pinene trials from day 1 to day 4 of training with the presence of odor landmarks. PV activity (combination of all recorded neurons in all 5 mice) generated from mean activity on all trials normalized to the max mean activity for each neuron. Trajectories were plotted in the dimensions of the first 2 principal components.

### A model of place cells driven by and interacting with path integrators

Model place cells receive inputs that are modulated by a set of locations estimated by path integration (figure 4a). All of the model place cells receive the same set of 100 spatially tuned inputs with firing rates *f*_*i*_(*x*_*i*_) for *i* = 1, 2, …, 100. Each function *f*_*i*_ is generated initially by a Gaussian random process (Gaussian noise low-passed filtered with a length constant of 0.5 m) and then held fixed. Each variable *x*_*i*_*(t)* is an independent noisy estimate of the location of the animal at time *t*, obtained by integrating a noisy estimate of the animal’s velocity with added white-noise fluctuations. Specifically, on each trial, velocities for these integrators are chosen from a Gaussian distribution around the true velocity of the animal (taken to be 0. 4 m/s) with a standard deviation of 0.1 m/s. In addition, white noise (Gaussian with standard deviation of 0.004 m/s for an integration step size of 0.025 s) was added to the integrated velocity. This causes each *x*_*i*_ to differ from the others on every trial and also to vary from trial to trial. As a result, the modulated inputs, *f*_*i*_*(x*_*i*_*(t))*, are also different and vary from trial to trial. These fluctuations increase as a function of *t* (as the animal moves along the virtual track) because the integrator-to-integrator and trial-to-trial variance of the location estimates increases as a function of the integration interval (figure 4b).

The input to place cell *a*, for *a* = 1, 2, …, 150, is Σ_*i*_ *w*_*ai*_ *f*_*i*_*(x*_*i*_*(t))*/|*f(x(t))*|, where the expression in the denominator is the norm of the vector with components *f*_*i*_*(x*_*i*_*(t))*, and *w*_*ai*_ is the weight of the input from integrator *i* to place cell *a*. A threshold is subtracted from this input, and the place cells firing rate, *r*_*a*_*(t)*, is determined by rectifying the result.

All place cells that are inactive on a given trial are subject to plasticity at their synapses from the integrator-modulated inputs. For each inactive place cell, we choose a time *t*_*a*_* for this plasticity to take place, simulating the effects of a dendritic plateau potential^30,31^. We denote the values of the path integrators at this time and on this trial by *x*_*i*_\**(t*_*a*_*). The result of this plateau is that the weights to model place cell *a* are set to *w*_*ai*_ = *f*_*i*_*(x*_*i*_\**(t*_*a*_*))/|*f(x*\**(t*_*a*_*))|, i.e. the input at time of the plateau. Following this plasticity, the input to place cell *a* is equal to the cosine of the angle between the vector *f(x*\**(t*_*a*_*)) (the input vector at time *t*\* on the trial when the plasticity occurred) and the vector *f(x)* at the current time on the current trial. If the place cell happened to form near the start of the virtual track, it is likely that it will fire on subsequent trials because the vector *f(x)* only fluctuates by a small amount from trial to trial when the integrators only have to integrate over a short distance. If, on the other hand, the place cell formed at a larger distance from the start, the larger fluctuations in *f(x)* from trial to trial cause a poor match to the weights and, as a result, the place cell is unlikely to fire. This is the reason that reliable place cells only form across the first 2 m of the virtual track (figure 4c).

Thus far, we have described the connections from the path integrators to the place cells, but there are connections from place cells to path integrators in the model as well (figure 4a), and these are also plastic. When plasticity acts on the inputs to place cell *a*, we imagine that it also acts on the inputs from that place cell back to the path integrators. This is assumed to be similar to the plasticity discussed in reference^7^, but we do not model this circuit in full, focusing instead on the results of this plasticity. The effect of this plasticity is that the value *x*\*_*i*_*(t*_*a*_*) is stored in synapses from place cell *a* to a path integrator *i*. Specifically, if an odor is present at time *t*_odor_, which we assume gates the effect of place cells on the path integrators (figure 4a), path integrator *i* is reset to *x*_*i*_*(t*_odor_) = Σ_*a*_ *x*\*_*i*_*(t*_*a*_\**)r*_*a*_*(t*_odor_)/Σ_*a*_ *r*_*a*_*(t*_odor_). The result of this resetting is that, after the odor appears, the trial-to-trial variability of the path integrator estimates is greatly reduced (figure 4d). This consistency produces a better match between the weight vectors of place cells formed beyond the odor location and the input vectors generated by the path integrators. The result is that reliable place cells can now form along the entire virtual track (figure 4e).

