## Supplemental Figures for "Olfactory Landmarks and Path Integration Converge to Form a Cognitive Spatial Map"

**Supplemental Figure 1**

**a,b,c,** Mean speed at positions along the track for different trial types (n = 5). **a,** Last day of training with no odor and on 4$^{th}$ day of training with odor. **b,** Training with no odor before and after training with odor landmarks. **c,** On each day of training with odor landmarks.

**Supplemental Figure 2**

**a,b,** Trial to trial variability as a function of distance from landmarks. Black, last session with no odor landmarks. Red, on 4th day with odor landmarks. **a,** Mean Pearson correlation of the activity population vector of all recorded cells in each mouse (n = 5) over distance bins comparing even numbered and odd numbered trials. **b,** Mean percent of trials with zscore > 1 in place field (peak bin +/- 10cm) for all recorded cells plotted according to location of peak mean activity.

**Supplemental Figure 3**

Mean Pearson correlation (n = 5, error bars = SEM) of the activity population vector between limonene and pinene trials over corresponding areas of the track (0-1m, 1-2m, 2-3m, and 3-4m).

**Supplemental Figure 4**

**a,** Activity of combined place cells for all mice (n = 5) along the virtual track on 1st day of training with odor landmarks (9.2%, 222/2405). **b,c,** Activity of combined odor cells (n = 5) along the virtual track. **b,** 1 day of training with odor landmarks (5.7%, 137/2405). **c,** 4 days of training with odor landmarks (2.4%, 68/2778)

**Supplemental Figure 5**

**a, b,** Population vector trajectory along the virtual track with odor landmarks in the 2-dimensional space defined by the first 2 principal components for all recorded cells. **a,** 2 days of training. **b,** 3 days of training.

**Supplemental figure 1**

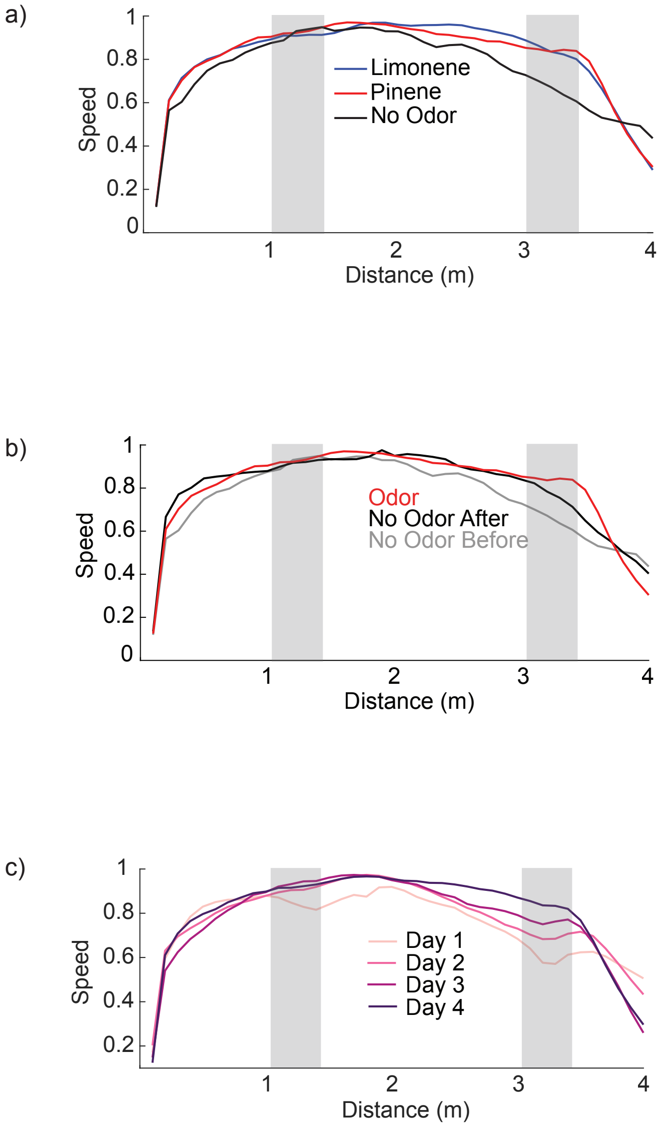

**Supplemental figure 2**

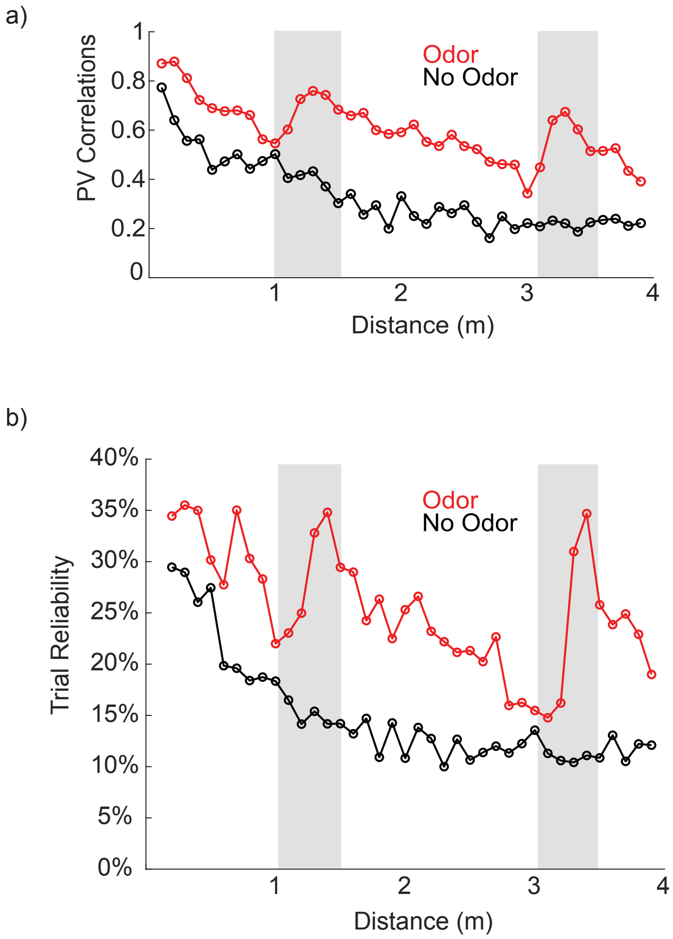

**Supplemental figure 3**

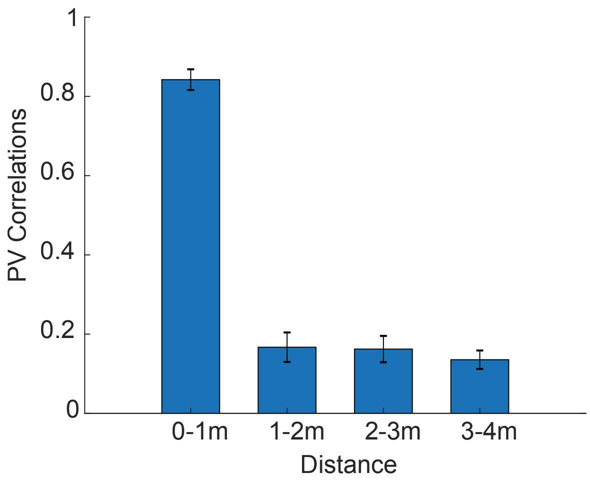

**Supplemental figure 4**

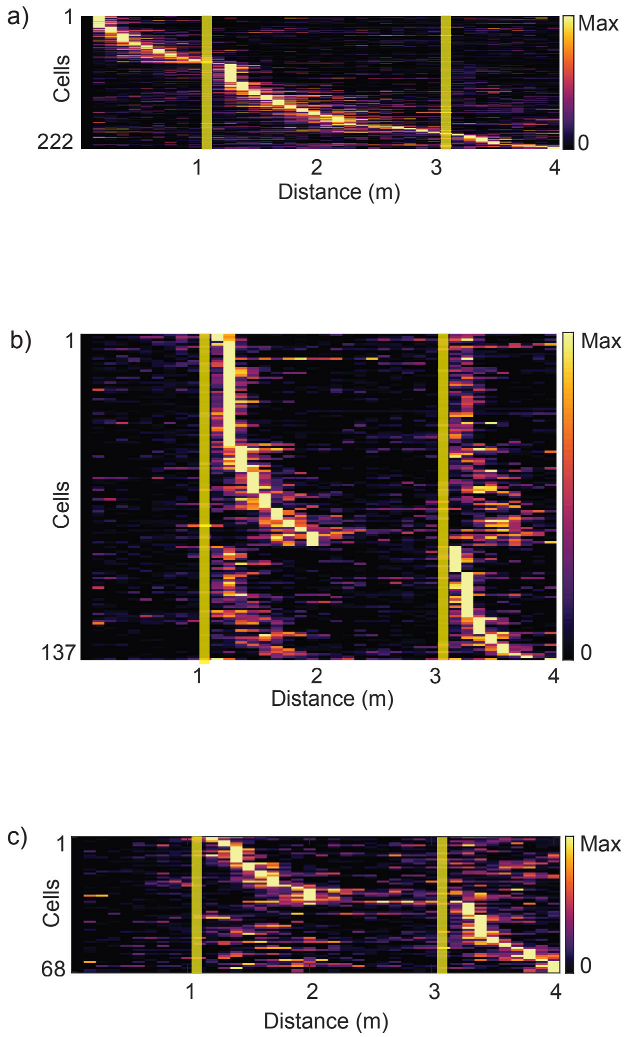

**Supplemental figure 5**

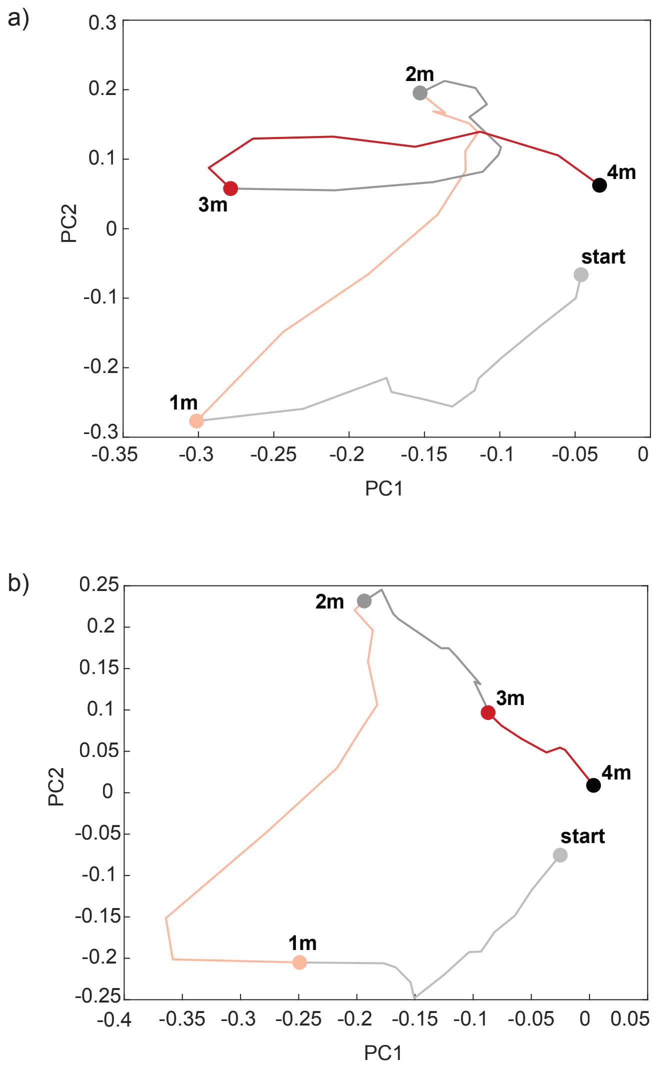
